# Fabrication and Modeling of Recessed Traces for Silicon-Based Neural Microelectrodes

**DOI:** 10.1101/2020.04.06.028159

**Authors:** Nicholas F Nolta, Pejman Ghelich, Martin Han

**Affiliations:** Department of Biomedical Engineering, University of Connecticut, Storrs, CT, USA; Institute of Materials Science, University of Connecticut, Storrs, CT, USA

**Keywords:** microelectrode, neural electrode, fabrication, insulation, planarization, mechanical stress

## Abstract

Chronically-implanted neural microelectrodes are powerful tools for neuroscience research and emerging clinical applications, but their usefulness is limited by their tendency to fail after months *in vivo*. One failure mode is the degradation of insulation materials that protect the conductive traces from the saline environment. Studies have shown that material degradation is accelerated by mechanical stresses, which tend to concentrate on raised topographies such as conducting traces. Therefore, to avoid raised topographies, we developed a fabrication technique that recesses (buries) the traces in dry-etched, self-aligned trenches. The depth of the trenches and the thickness of the traces are matched so that overlying insulation materials are flat, which, according to finite-element modeling, reduces stress concentrations in the insulation material. Here, we provide details on process optimization, modeling of intrinsic stress, and characterization using SEM, focused ion-beam cross sections, profilometry, and electrochemical impedance testing. The technique requires no extra masks, is easy to integrate with existing processes, and produces flatness within about 10 nm.

## 1. Introduction

Clinical deep brain stimulators [1], cochlear implants [2, 3], and spinal cord stimulators [4, 5] are used for treatment of Parkinson’s disease, essential tremor, depression, obsessive-compulsive disorder, obesity, deafness, and chronic pain in cases where other options have failed. Collectively, these systems have benefitted hundreds of thousands of patients and demonstrated the signficant clinical potential of interfacing with the nervous system even at a relatively crude level.

Meanwhile, in research labs, the development of smaller, more complex devices has allowed for higher-bandwidth and more intimate contact with the nervous system. Since 1970 [6], many of these devices have been produced using microelectromechanical systems (MEMS) fabrication technology, enabling studies of next-generation deep brain stimulators [7–9] and increasingly sophisticated visual [10, 11], auditory [12–14], and tactile/motor [15, 16] prostheses in animals and humans.

Unfortunately, advanced functionality has come at a cost to reliability. While there are examples of microelectrodes performing successfully for many years, in most cases, performance declines over months [17–19]. Understanding the causes of microelectrode failure is an active area of research. The prevailing view is that a variety of factors co-contribute, including device materials degradation, encapsulation by glia, changes in neuronal health/activity/connectivity, and gross movement of the device over time [18–24]. In order to bring advanced neural microelectrodes into the clinic, engineers will need to mitigate each of these failure modes [25].

Materials degradation has been studied for a variety of device types and materials. Most commonly, the insulation materials develop cracks or pinholes [19–21, 26–28], exposing too much of the electrode sites [20, 21, 28], creating short-circuits and crosstalk [27, 29, 30], or causing electrode site material to detach [28, 31]. Both ceramics [19, 31] and polymers [20, 21, 26–30] are vulnerable to degradation and no material has yet established itself as immune to degradation.

While most studies focused on chemical mechanisms, relatively few have examined the effects of mechanical stress on neural electrode materials failure. A study by Schmitt et al. compared the reliability of several insulation strategies in saline solution [32]. The authors used a wet etch to fabricate devices with recessed (buried) traces, and found that these devices lasted five times longer than non-recessed devices with identical insulation material. The authors noticed delamination of insulation materials over non-recessed traces and theorized that the raised topography of the non-recessed traces concentrated intrinsic stresses generated during fabrication, and accelerated materials degradation through a process called stress corrosion cracking. A different study by Kozai et al. looked at material failures in NeuroNexus (Ann Arbor, MI, USA) silicon microelectrodes implanted in mice for 19-27 weeks [19]. The authors reported cracking and corrosion of insulation materials over the traces, especially near the iridium electrode sites. Mechanical modeling of extrinsic stresses (from device movement relative to the brain) showed that these areas also experienced up to three times higher stresses due to the combined effects of raised traces and proximity to stiff iridium electrode sites. Although these issues may be specific to the design of NeuroNexus electrodes, it nevertheless supports the idea that mechanical stress may influence the longevity of insulation materials.

Here, we present a fabrication technique similar to Schmitt et al., but employing a dry etch, which is easier to implement. We successfully incorporated the technique into our production process for functional probes, similar to those we have used extensively in long-term animal studies [12, 13]. We previously described our method in a short conference publication [33]. In this report, we provide greater detail on the method and its optimization and use finite element modeling to show its predicted effect on intrinsic stress.

## 2. Methods

### 2.1. Wafers and photolithography

As shown in figure 1(*a*), the process began with four-inch-diameter silicon-on-insulator wafers with 1 μm wet-grown thermal oxide on both sides (Ultrasil, Hayward, CA, USA). These wafers had a ‘device’ silicon layer 80 μm thick, a ‘buried oxide’ thermal oxide layer 2 μm thick, and a ‘handle’ silicon layer 350 μm thick. Silicon-on-insulator wafers were used for functional devices, while regular oxidized silicon wafers were used for test samples. In figure 1(*b1*), AZ 5214E-IR photoresist from Merck (Darmstadt, Germany) was patterned using a standard image reversal lithography protocol. Traces were 10 μm wide and 10 μm apart, with 5 μm wide curved features around electrode sites.

**Figure 1.**
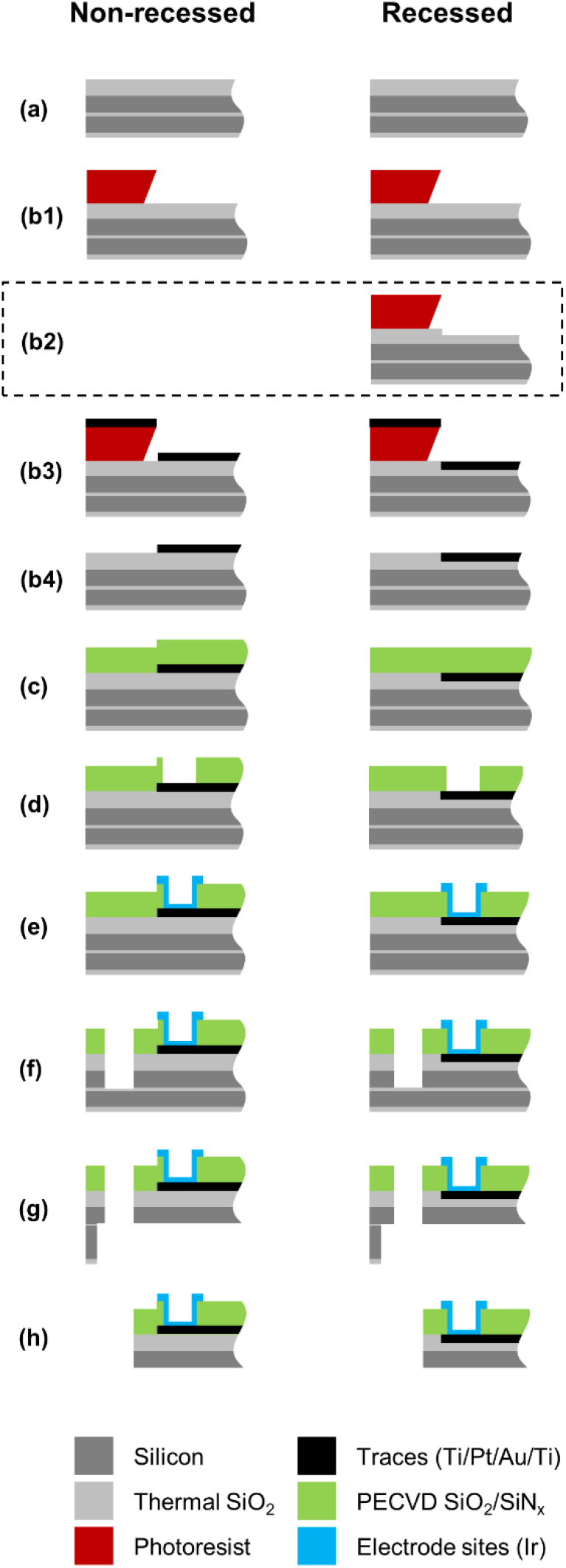
Overview of the fabrication process with an emphasis on the electrode site. A side-view cross section near the tip of a device is depicted; the rest of the shank and the bond pad areas are off to the right and not shown. Not to scale. (*a*) Initial oxidized silicon-on-insulator wafer. (*b1*-*b4*) Recessed traces process, including (*b1*) photolithography, (*b2*) reactive ion etching (recessed traces only), (*b3*) trace metal deposition, and (*b4*) lift-off and plasma cleaning. This was followed by (*c*) deposition of insulation layers, (*d*) opening of vias, (*e*) sputter deposition of electrode site metal, (*f*) etching the shape of the shanks, (*g*) backside thinning, and (*h*) manual release. The critical difference in the two processes is the recessed traces etch step *b2* which results in trace metal positioned inside the thermal SiO_2_.

### 2.2. Reactive ion etching to pattern trenches

This step is depicted in figure 1(*b2*). Wafers were etched in an NLD-570 inductively-coupled plasma RIE (ULVAC Technologies, Inc., Methuen, MA, USA) with 90 sscm argon, 10 sscm octafluoropropane, 3 mTorr pressure, 1 100 W coil power, and 227 W platen power, at 5 °C. The chamber was O_2_-cleaned for 20 min and the etch was run for 5 min on a blank wafer as preconditioning before etching. This recipe had an etch rate of 410 nm min^−1^ and a selectivity of approximately 2:1 for silicon dioxide vs. photoresist.

### 2.3. Metal deposition

As shown in figure 1(*b3*), an electron beam evaporator (Denton Vacuum, LLC, Moorestown, NJ, USA) with throw distance approximately 50 cm was used to deposit a metal stack of 30 nm titanium, 20 nm platinum, 300 nm gold, and optionally an additional 50 nm titanium at rates of 1.0, 1.5, 2.0, and 1.0 Å s^−1^, respectively. Samples were positioned in the center, directly facing the source, with rotation turned on. For functional devices, a metal stack of 30 nm titanium, 20 nm platinum, and 300 nm gold was subsequently deposited on the bond pads only.

### 2.4. Lift-off, cleaning, and argon etching

As depicted in figure 1(*b4*), samples were immersed in acetone for at least 1.5 h. The samples were then transferred to fresh acetone, sonicated for 5 min, and rinsed with isopropanol and water. Next, an oxygen plasma barrel asher (SCE 106, Anatech Ltd., Sparks, NV, USA) was used to remove remaining polymeric residues for 5 min at 150 W, 40 sscm, and 435 mTorr. An optional argon etch in the ULVAC NLD-570 ICP RIE was performed at 5 °C, 3 mTorr, 600 W coil, 145 W platen, 100 sscm argon to improve yield of devices near the edges of the wafer. Finally, less than 30 min before deposition of insulation layers, samples were cleaned by dipping into a solution of 100 ml water, 20 ml 29 % NH_4_OH, and 20 ml 30 % H_2_O_2_ for 5 s then rinsed with water.

### 2.5. Insulation

As shown in figure 1(*c*), eight layers of silicon nitride (N) and silicon dioxide (O) were deposited in the order NONONONO to a total thickness of 1.85 μm using a parallel electrode PECVD reactor (SPTS Technologies Ltd, Newport, UK). The recipes used 300 °C, 900 mTorr, and the high frequency generator at 20 W for silicon nitride and 30 W for silicon dioxide. The gas flows for silicon nitride were 35 sscm SiH_4_, 55 sscm NH_3_, and 1 960 sscm N_2_; for silicon dioxide, 10 sscm SiH_4_, 1 420 sscm N_2_O, and 392 sscm N_2_.

### 2.6. Functional device fabrication

Figure 1(*d*) shows the etching of vias by RIE, which opened the electrode sites (shown) and bond pads (not shown). After this, electrode site metal was deposited by sputtering titanium at 200 W DC for 8 min 20 s (approximately 50 nm) followed by iridium at 100 W DC for 30 min (approximately 200 nm) with rotation on (figure 1(*e*)). The pressure was 4 mTorr and gas flow was 15 sscm argon. Next, the shapes of the shanks were defined by first using RIE to etch through the PECVD insulation and the thermal SiO_2_. Then, Bosch-process deep reactive ion etching (DRIE) was performed in a Rapier DRIE (SPTS Technologies Ltd, Newport, UK) (figure 1(*f*)). The buried oxide in the silicon-on-insulator wafer served as the etch stop. To free the devices from the handle wafer, the 4” wafers were adhered to a 6” oxidized silicon wafer using a thermally-conductive paste (Cool-Grease, AI Technology, Princeton Junction, NJ, USA) then etched from the backside using an SF6-based etch in the Rapier DRIE for approximately 30 minutes (figure 1(*g*)). The buried oxide was also etched from the backside using RIE. Finally, the devices were released using forceps (figure 1(*h*)). This process represents the current evolution of a process initially developed by Han et al. [13]. Finished microelectrodes were then epoxied onto custom printed circuit boards, wire bonded, and over-coated with Epotek 301 epoxy (Epoxy Technologies, Inc., Billerica, MA, USA).

### 2.7. Characterization

Scanning electron microscopy (SEM) images were collected using 2-5 kV accelerating voltage and either the in-lens or in-chamber secondary electron detectors in a Hitachi (Chiyoda, Tokyo, Japan) SU8230 SEM. Profilometry was performed with a DektakXT stylus profilometer (Bruker, Inc., Billerica, MA, USA) with 12.5 μm stylus tip. Focused ion beam (FIB) cross section analysis was performed using a ZEISS (Oberkochen, Germany) Crossbeam with gallium ion source. An Autolab PGSTAT128N potentiostat/galvanostat (Metrohm AG, Herisau, Switzerland) was used to acquire AC electrochemical impedance measurements in room-temperature phosphate buffered saline, pH 7.4, using a large platinum counter electrode, an Ag/AgCl reference electrode, and a 10 mV peak-to-peak sinusoid at 1 kHz.

### 2.8. Finite element modeling

Finite element modeling was carried out in COMSOL’s structural mechanics module (COMSOL, Inc., Burlington, MA, USA). 2D cross sections of shanks with non-recessed or recessed traces were drawn with accurate dimensions. The model assumed isotropic material properties using the values for silicon, silicon dioxide, silicon nitride, and gold from COMSOL’s MEMS material library. Based on characterization data, the PECVD silicon dioxide was given −140 MPa (compressive) intrinsic stress and PECVD silicon nitride +330 MPa (tensile) intrinsic stress. Based on a literature value [34], the thermally-grown silicon dioxide was assumed to have −300 MPa (compressive) intrinsic stress. The outer boundaries were free and no external forces were applied.

## 3. Results

### 3.1. Characterization

Figure 2(*a*) shows an optical image of a non-recessed device after insulation, via etching, electrode site metallization, and shank shaping. The traces are visible through the transparent insulation. To quantify the degree of flatness, stylus profilometry was performed on both types of devices. We found that flatness within about 10 nm could be obtained for recessed traces, in contrast to non-recessed traces which were raised about 350 nm (figure 2(*b*)). After backside etching and release, the devices were examined using SEM. At low magnification, the raised topography of insulation covering non-recessed traces was easily visible, while the recessed traces were flat enough as to be invisible (figure 2(*c*)). To understand the interior structure of the traces, high-magnification FIB cross sections were obtained (figure 2(*d*)). The faint contrast between the repeating layers of silicon dioxide and silicon nitride insulation revealed how the insulation curves sharply upward over non-recessed traces but is nearly flat over recessed traces. These images also confirmed the plausibility of our finite element model geometry (see section 3.2).

**Figure 2.**
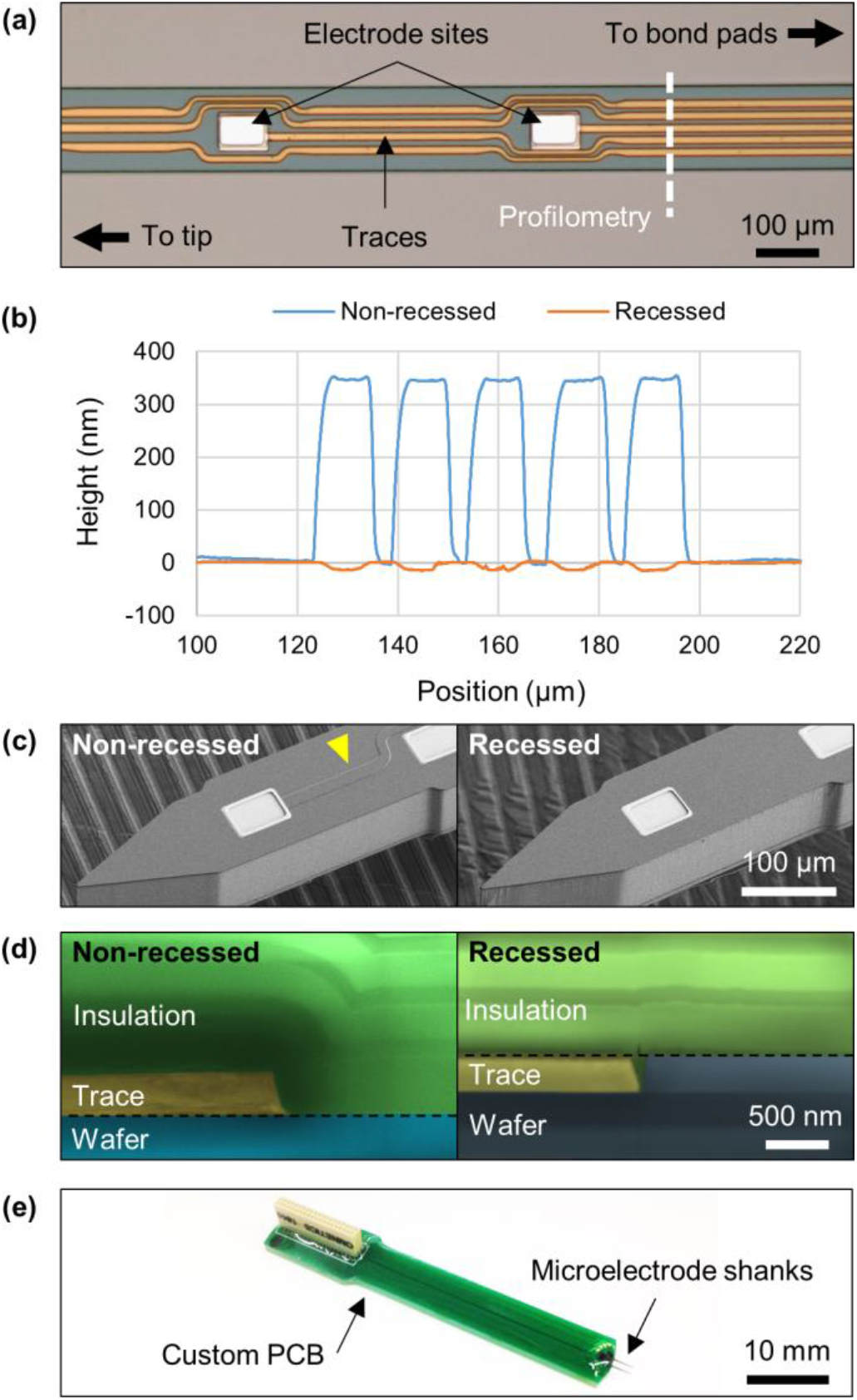
(*a*) Light micrograph of a completed non-recessed device. The middle of a shank is shown, with the tip to the left (not shown) and bond pads to the right (not shown). The traces are visible through the optically-transparent insulation. White dotted line indicates the path along which the profilometer measured the heights of the traces. (*b*) Stylus profilometry across an area with five traces running parallel to one another for a non-recessed device and a recessed device, both after passivation. The non-recessed traces are 350 nm tall, corresponding to the 350 nm thick metal, whereas the recessed traces are flush with the wafer surface within about 10 nm. Note that height is exaggerated about 100-fold for visibility. (*c*) Completed, released microelectrodes imaged by SEM. The raised topography of the non-recessed traces (left) is easily visualized (yellow arrowhead), while the recessed traces are nearly invisible at this magnification. (*d*) FIB cross section revealing the internal structure of a non-recessed (left) and recessed trace (right). The image has been false-colored to assist in interpretation. The insulation layers curve sharply up and over the non-recessed trace, but are nearly flat over the recessed trace. Dotted line indicates wafer surface. (*e*) A two-shank microelectrode array wire-bonded and epoxied onto a custom printed circuit board for electrochemical testing.

After metal deposition and lift-off, small metal ‘fences’ were found towards the outer 50 % of the wafer, and only on specific sides of the features corresponding to the direction of deposition. The probable mechanism of fence formation is illustrated in figure 3. In an electron beam evaporator, metal is evaporated in a crucible some distance (here, about 50 cm) away from the wafer. The evaporated metal then showers onto the wafer in nearly straight line trajectories. Since the spot where the metal is evaporated by an electron beam is very small, the metal essentially comes from a point source 50 cm from the wafer. Therefore, in the center, the trajectory is nearly vertical, and in this area no fences were formed (figure 3(*a*-*c*)). Meanwhile, towards the edges, the trajectory is slightly angled, creating a widened gap on one side and fences on the other side (figure 3(*d*-*f*)). The angle would have been 2.9° at 1” from the center and 5.7° at the edge of the wafer. SEM inspection of a recessed trace immediately after metal deposition revealed that the fences formed without actually bridging the photoresist’s undercut (figure 3(*g*)). This rules out the possibility that fences were due to poor lift-off, which can happen if photoresist thickness and/or undercut is insufficient (causing metal to bridge the gap and create torn-off structures when physically separated).

**Figure 3.**
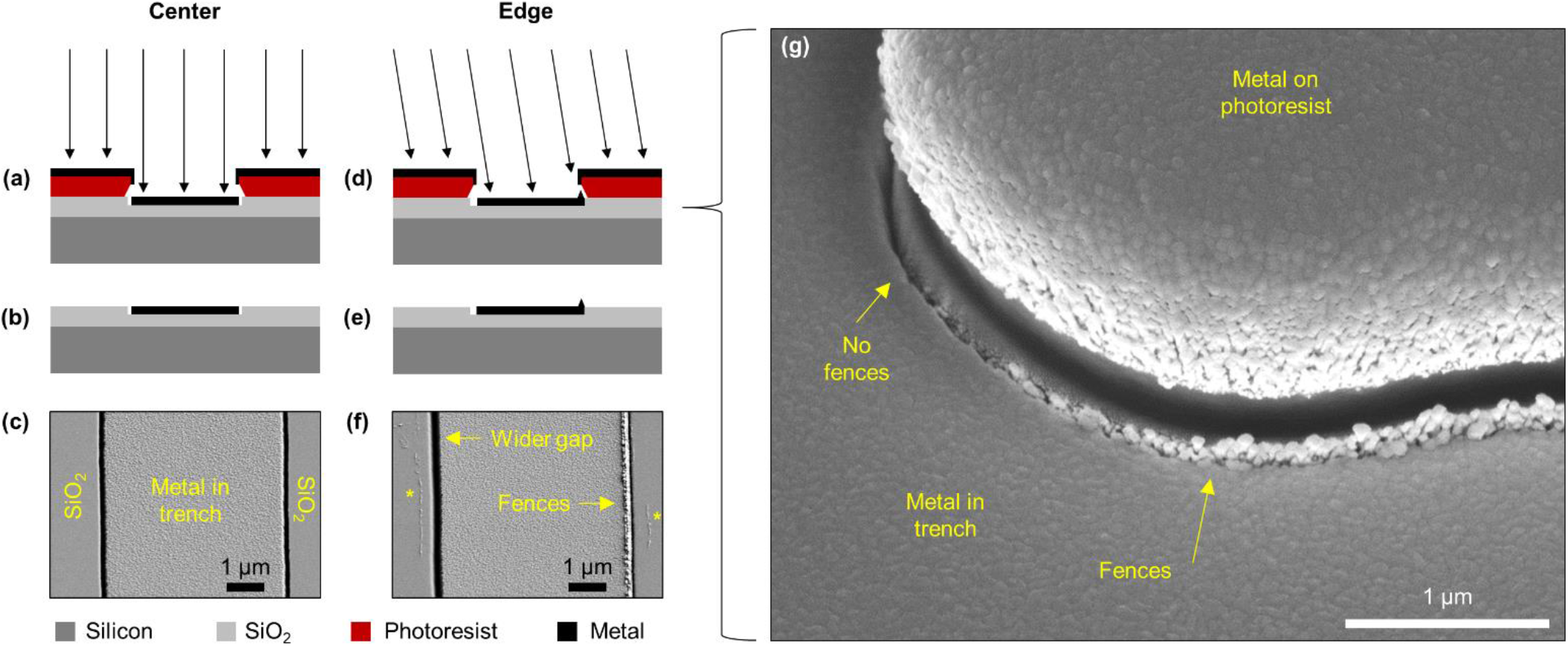
Likely mechanism of fence formation. (*a*) In the center of the wafer, metal is deposited vertically and lands in the middle of the trench. (*b*) Lift-off removes metal everywhere besides the trench. (*c*) After lift-off, SEM shows the metal was squarely centered in the trench and had no fences. (*d*) In contrast, towards the edges of the wafer, metal is deposited at a slight angle, resulting in a widened gap on one side and fences on the other. (*e*) After lift-off, the fences remain. (*f*) SEM image of a trace near the edge of the wafer. Asterisks denote polymer residues, which were present alongside traces all over the wafer, but were successfully cleaned off by oxygen plasma. (*g*) A recessed trace near the edge of the wafer immediately after metal deposition. This 90° corner feature reveals the direction-dependent nature of the fences: fences appeared only along edges corresponding to the direction of metal deposition, and did not connect to the metal on top of the photoresist.

Finally, to determine whether recessed traces interfered negatively with device performance, functional devices (figure 2(*e*)) were tested in PBS. AC impedances at 1 kHz were 97 ± 9 kΩ, which is typical of devices with sputtered iridium microelectrodes of this size (2 000 μm^2^). This benchtop measurement was not indented to show improved long-term performance—only that recessed traces do not interfere with normal functioning of the electrode.

### 3.2. Finite element modeling

To confirm whether recessed traces could be expected to reduce stress concentrations, finite element modeling was performed. Figure 4 shows von Mises stress in the vicinity of a non-recessed and a recessed trace. For the non-recessed trace, each of the interior corners of the PECVD silicon dioxide/silicon nitride layers show much higher stress than the surrounding areas. In the recessed trace, these stress concentrations are absent, however the stress does change slightly (higher for silicon dioxide, lower for silicon nitride) over the metal trace, most likely due to the change in thickness of thermal silicon dioxide. Some high stresses are also present in the lowermost SiN_x_ insulation layer at the boundary of the metal trace.

**Figure 4.**
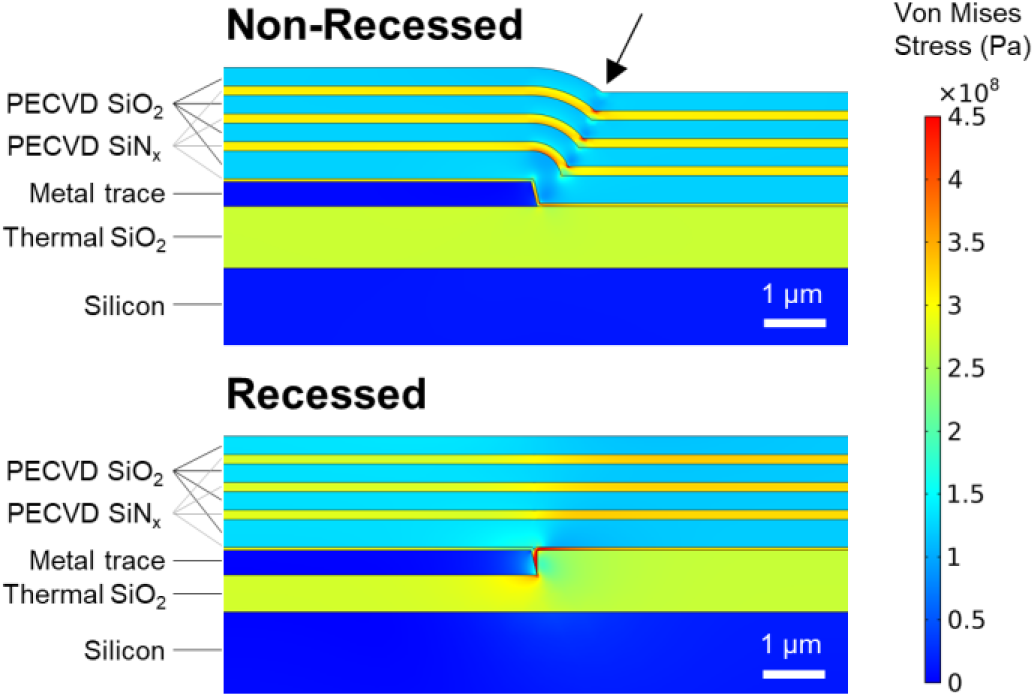
Von Mises stress near a non-recessed (top) and a recessed trace (bottom), with geometry similar to the actual device profiles shown in figure 2. For the non-recessed trace, the interior corners of the PECVD silicon dioxide/silicon layers (arrow) show higher stress than the surrounding areas.

## 4. Discussion

Overall, we have demonstrated a simple technique that allows traces to be recessed (buried) within a substrate. The technique does not require any additional masks, alignment, or polishing steps, only dry etching steps, and does not interfere with the functionality of the device. Our technique is potentially applicable to other neural microelectrode designs where the conducting traces are sandwiched between two insulating layers. Recessing the electrode sites (not just the traces) could have the additional benefit of reducing the edge effect of charge accumulation during stimulation.

We believe that the degree of flatness achieved with this technique is more than adequate for the purposes of reducing stress concentrations. Insulation layers are typically at least 1 μm thick, so 10 nm represents only 1 % of their total thickness. The remaining source of error is mainly due to the variability of the etch step. If better planarity is desired, the technique could be improved by measuring the depth of the trench after the etch and then making fine adjustments to the deposition thickness. Alternatively, a slow oxide etch, metal etch, or argon etch could be used to make corrections after metal deposition.

The most significant limitation of this technique in its current form, in our opinion, is yield. Perfect, fence-free traces could only be found in the center 2” of the 4” wafer, which is acceptable for research purposes, but would not be ideal for industrial scale-up. We have several ideas for improvement. Increasing the throw distance (distance between metal crucible and wafer) would help by making the angle of deposition more vertical. Preserving more photoresist undercut would also help, though this would also create a slightly larger gap between metal and trench sidewall. Another tactic would be to try to remove the fences after they form. We attempted this by swabbing and sonicating in acetone and using an argon RIE, but these were only partly successful. High-pressure solvent cleaning or aggressive scrubbing could be explored.

Finite element modeling showed that non-recessed traces are likely to create intrinsic stress concentrations that recessed traces do not. Intrinsic stresses are formed due to the different coefficients of thermal expansion of different materials when they are deposited at high temperatures then cooled to room temperature. Schmitt et al. did not model intrinsic stresses, so our findings add new support to their hypothesis that reduced concentrations of intrinsic stresses could be responsible for the improved performance of recessed traces in their study [32]. Our model’s accuracy could be improved with more accurate geometry, more precise estimates of intrinsic stresses, and a more detailed representation of the metal layers. We did not model extrinsic stresses (stresses due to externally-applied forces) because this aspect was already explored in the study by Kozai et al. which showed that raised, non-recessed traces tended to concentrate extrinsic stresses [19].

An alternative to our technique is chemical-mechanical polishing (CMP), which is the industry standard for planarization (used for example in the popular dual-Damscene process). Our technique’s advantages over CMP are its simplicity and its availability to cleanrooms lacking CMP equipment. To accomplish the same effect using CMP, insulation would need to be deposited at least as thick as the trace. Then, CMP would remove material until the raised topographies over the traces were fully eroded. Finally, additional insulation would be deposited on top of the planar surface.

The next step in this work will be to test whether our recessed traces withstand degradation better than non-recessed traces in an accelerated soak test. If they do, our technique represents a simple and broadly-applicable approach for improving the reliability of neural interfaces and MEMS in general [35].

## 5. Conclusion

A technique for recessing the conducting traces of a neural probe within the substrate in order to reduce stress concentrations was described in detail. The technique requires no extra masks and allows the insulation layers to be flat within about 10 nm. The improved flatness is expected to increase the chronic reliability of neural electrodes, a hypothesis that will be tested in future *in vitro* and *in vivo* studies.

## Acknowledgements

This work was supported by the NIH research grants R01DC014044 and R24NS086603 (MH). We would like to thank Joseph Favata and Sina Shahbazmohamadi at the University of Connecticut for performing FIB analysis and Ling Xie and Kenlin Huang at the Center for Nanoscale Systems at Harvard University for help with the RIE recipes.

